## Supplementary for "Metabolic robustness to growth temperature of a cold adapted bacterium"

**Table S1.** Main features of the transcriptomic dataset BioProject **PRJNA886636.** Reads numbers refer to the raw, unfiltered files.

| **Accession Number** | **Number of Reads** | **Temperature** |
| --- | --- | --- |
| SRX17780709 | 18,376,681 | 15°C |
| SRX17780711 | 25,089,797 | 15°C |
| SRX17780712 | 31,526,393 | 15°C |
| SRX17780713 | 17,586,523 | 0°C |
| SRX17780714 | 23,037,026 | 0°C |
| SRX17780710 | 29,173,509 | 0°C |

**Table S2.** Expression data for DNA-replication related-genes

| **Locus tag** | **log_2_FC** | **Adj. p-value** | **Annotation** |
| --- | --- | --- | --- |
| PSHAa0090 | 1.1108113 | 0.00000925 | \|xerC\| site-specific recombinase |
| PSHAa1218 | 1.8251366 | 1.52E-13 | \|recN\| DNA repair protein recN |
| PSHAa2356 | 1.0226152 | 0.0000609 | \|topB\| DNA topoisomerase III |
| PSHAa2361 | 1.2276516 | 0.0000048 | DNA topoisomerase III (N terminal part) |
| PSHAa2873 | 1.1514558 | 0.000014 | \|lexA\| transcriptional repressor for SOS response |
| PSHA_RS17095 | 1.382628818 | 0.00000669 | \|xni\| putative exonuclease IX |

­

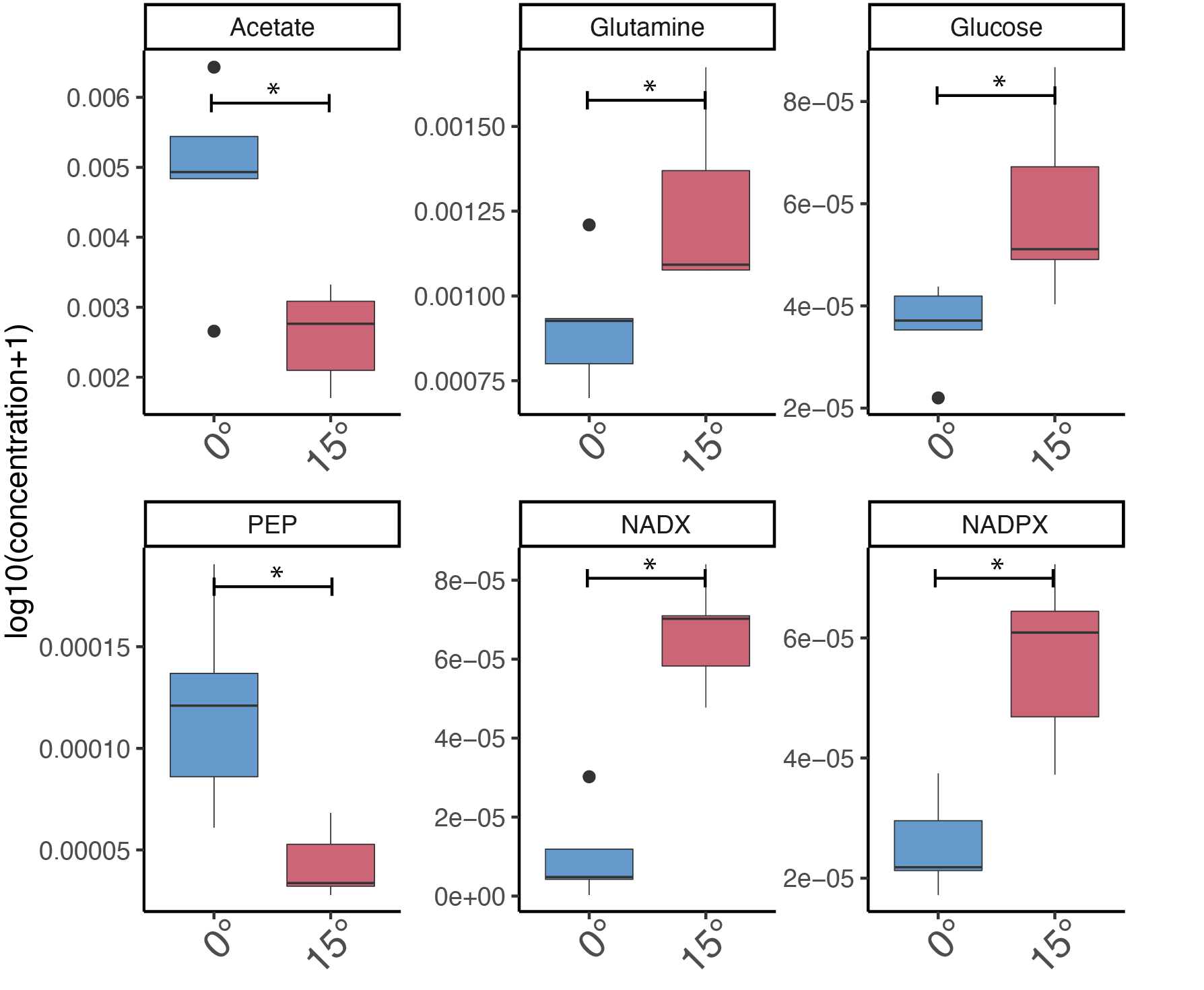

**Figure S1.** Boxplots of intracellular metabolites that showed a statistically significant difference concentration across all the time points analysed.

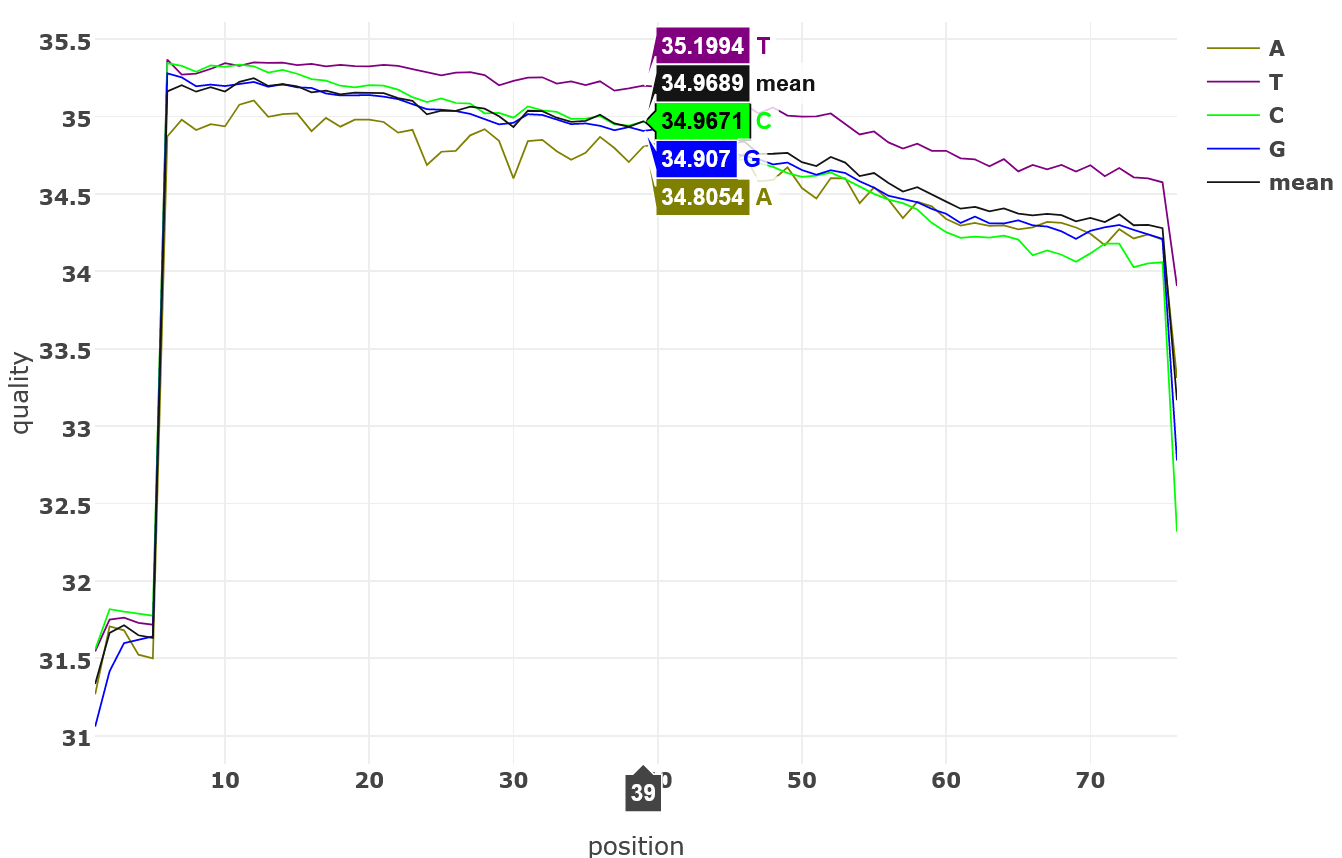

**Figure S2.** Quality check report performed using fastp (Shifu Chen et al., 2018). Position 39 is being displayed as a sample representative of the quality at read mid-length.

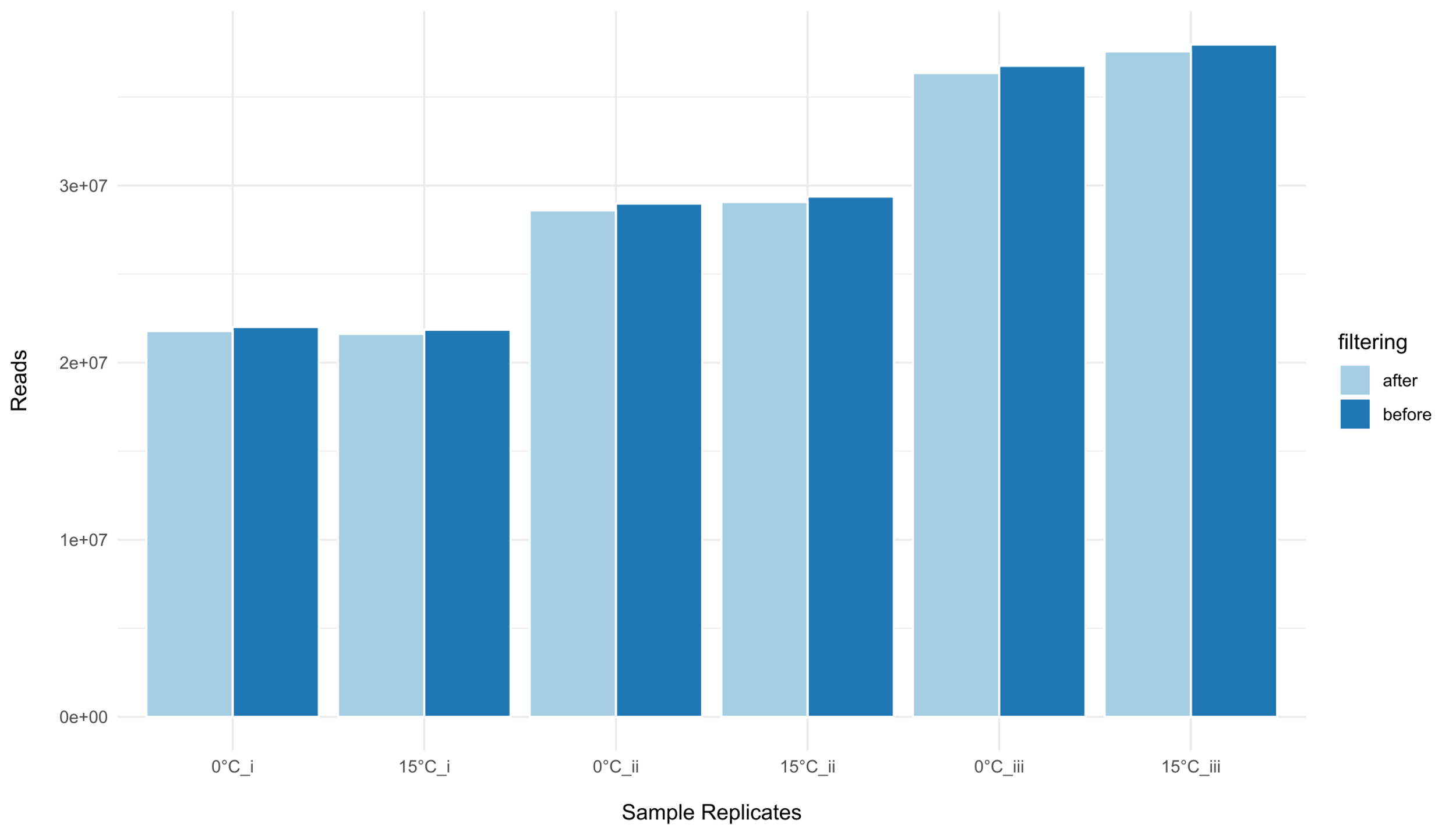

**Figure S3.** Sample replicates reads count before and after QC with fastp.

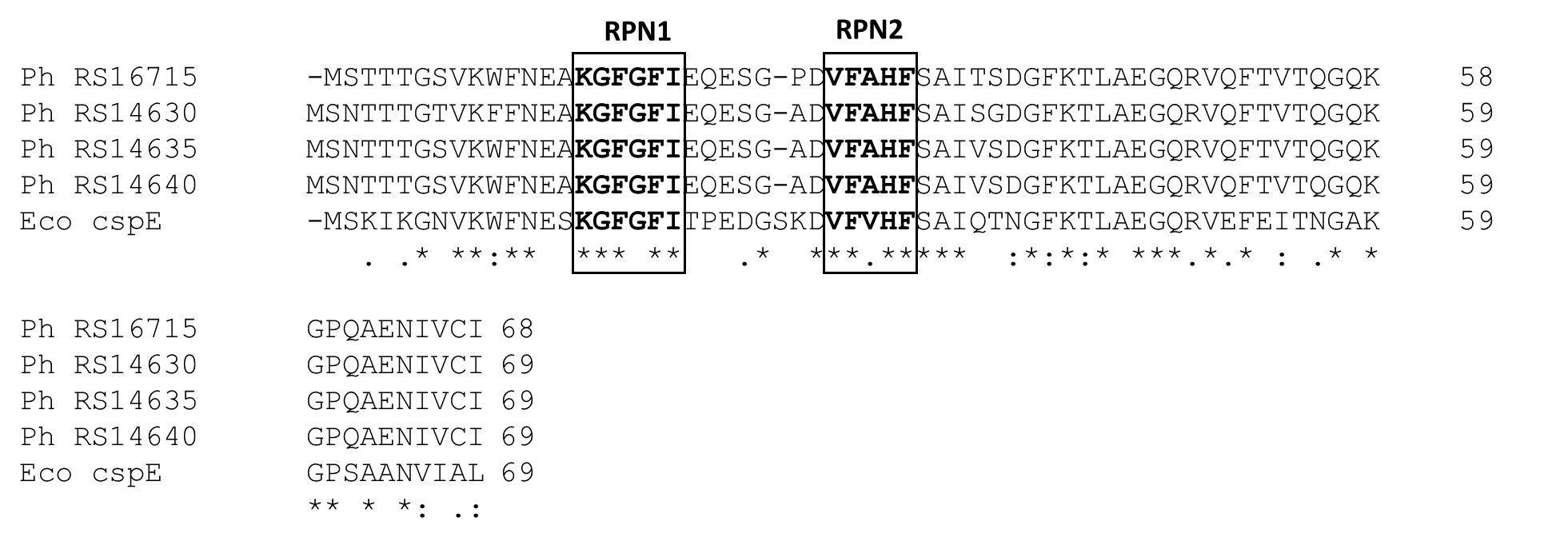

**Figure S4.** Sequence alignment of four psychrophilic CSPs with the mesophilic CspE protein. Gaps indicated by hyphens were introduced to improve alignment, and identical amino acids are indicated with asterisks. The RNA-binding motifs RNP1 and RNP2 are boxed in black.
